# Ripasudil in a model of pigmentary glaucoma

**DOI:** 10.1101/2020.05.04.076760

**Authors:** C Wang, Y Dang, S Waxman, Y Hong, P Shah, RT Loewen, X Xia, NA Loewen

## Abstract

**Purpose:** To investigate the effects of Ripasudil (K-115), a Rho-kinase inhibitor, in a porcine model of pigmentary glaucoma.

**Methods:** Hallmark features of trabecular meshwork (TM), the principle structure of the outflow system affected in this model, were analyzed. In vitro TM cells and ex vivo perfused eyes were subjected to pigment dispersion followed by K-115 treatment (PK115). PK115 was compared to sham-treated controls (C) and pigment (P). Cytoskeletal alterations were assessed by F-actin labeling. TM cell phagocytosis of fluorescent targets was evaluated by flow cytometry. Cell migration was studied with a wound-healing assay. Intraocular pressure was continuously monitored and compared to after the establishment of the pigmentary glaucoma model and after treatment with K-115.

**Results:** In vitro, the percentage of cells with stress fibers increased in response to pigment but declined sharply after treatment with K-115 (P: 32.8 +/- 2.9%; PK115: 11.6 +/- 3.3%, P < 0.001). Phagocytosis first declined but recovered after K-115 (P: 25.7+/-2.1%, PK115: 33.4+/-0.8%, P <0.01). Migration recuperated at 12h with K-115 treatment (P: 19.1+/-4.6 cells/high-power field, PK115: 42.5+/-1.6 cells/high-power field, P <0.001). Ex vivo, eyes became hypertensive from pigment dispersion but were normotensive after treatment with K-115 (P: 20.3 +/- 1.2 mmHg, PK115: 8.9 +/- 1.7 mmHg; P< 0.005).

**Conclusion:** In vitro, K-115 reduced TM stress fibers, restored phagocytosis, and restored migration of TM cells. Ex vivo, K-115 normalized intraocular pressure.

## Introduction

Pigment dispersion syndrome (PDS) is characterized by the shedding of pigmented debris from the iris pigment epithelium and its ectopic accumulation throughout the anterior segment of the eye but primarily in the trabecular meshwork (TM) [1]. Pigmentary glaucoma, a common secondary open-angle glaucoma, can develop due to PDS. It is characterized by heavy pigmentation of the TM, increased intraocular pressure (IOP), and characteristic glaucomatous optic neuropathy [2]. An increased outflow resistance results from an altered TM cytoskeleton, a decline in phagocytosis and migration, and activation of the Rho signaling pathway as can be shown both in vitro [3] and ex vivo [4].

Beta-blockers and prostaglandins effectively lower IOP in pigmentary glaucoma [5, 6]. Miotics lower IOP and reduce the underlying pigment dispersion by minimizing friction between zonules and the iris. However, this is poorly tolerated in non-presbyopic patients due to myopization [7]. Rho-kinase inhibitors are a more recent class of glaucoma medications that increase the outflow facility by relaxing the TM cytoskeleton [8–11] and dilating the distal outflow tract [12]. The cytoskeleton plays an important role in TM-mediated regulation of AH outflow. Ripasudil (K-115) inhibits both isoforms of rho-associated, coiled-coil-containing protein kinase (ROCK), ROCK-1, and ROCK-2 [13]. K-115 is an isoquinoline monohydrochloride dihydrate that binds to the ATP-dependent kinase domain in the hydrophobic cleft between the N- and C-terminal lobes of the kinase domain [14], just like fasudil [15] from which it is derived. It has an IC_50_ for ROCK 1 and ROCK 2 of 51 and 19 nM, respectively. Tanihara et al. found that a 0.4% solution of K-115 lowered the IOP by around 20% from a low baseline of 19 mmHg [16].

In this study, we hypothesized that K-115 can reverse pathological changes of the TM induced by pigment dispersion.

## Materials and Methods

### Generation of pigment

The generation of pigment followed our previous protocol [3]. Briefly, 10 pig eyes were obtained from a local meat market (Thoma Meat Market, Saxonburg, PA) within two hours of sacrifice. Irises were collected and frozen at −80 °C for 2 hours and thawed at room temperature. Freeze-thaw was repeated twice. Irises were added to 15mL of PBS, agitated to produce pigment particles, and filtered through a 70-μm cell strainer (431751, Corning Incorporated, Durham, NC). Particles were washed three times and suspended in 4mL of PBS as a stock solution.

### Pigmentary glaucoma model and K-115 treatment

The pigmentary glaucoma model was established as previously described [3]. Briefly, 24 paired pig eyes were obtained from a local meat market within two hours of sacrifice. The extraocular tissues were removed carefully. After disinfecting, the iris, lens, ciliary body, and posterior segment were removed. Anterior segments were mounted in perfusion dishes and perfused with Dulbecco’s modified Eagle’s medium supplemented with 1% fetal bovine serum (FBS, 10438026, Thermo Fisher Scientific, Waltham, MA) and 1% antibiotics (15240062, Thermo Fisher Scientific, Waltham, MA) at 3 μl/min. IOPs were recorded every 2 minutes with transducers (SP844; MEMSCAP, Skoppum, Norway) using LabChart software (ADInstruments, Colorado Springs, CO, USA). After 48h, the media was supplemented with either 1.67 × 10^7^ particles/ml of pigment granules (P, n=8), pigment plus 10 μM K-115 (PK115, n=7), or vehicle control consisting of normal perfusion medium (C, n=8).

### Actin cytoskeleton staining of primary TM cells

We used primary TM cells that were obtained and characterized by assays as described previously[3]. For all in vitro experiments, TM cells were seeded into six-well plates at a concentration of 1 × 10^5^ cells/well. Pigment media contained 1.67 × 10^7^ particles/ml. Stress fiber formation was evaluated through the labeling of F-actin. TM cells were seeded into one six-well plate with a coverglass slip placed at the bottom of each well. After reaching 80% confluency, six wells were treated with pigment media, and three wells were treated with vehicle control. After 24 hours, three of the pigment treated wells were additionally treated with 10 μM K-115 for 24h. Cells were fixed with 4% PFA for one hour at room temperature and washed with PBS three times. Samples were incubated with Alexa Fluor 488 Phalloidin (1:60 dilution, A12379, Thermo Fisher, Waltham, MA) for 30 min. DAPI was used to stain cell nuclei. 10 images of each group were acquired with an upright laser scanning confocal microscope (info) at 600x magnification. The percentage of cells with stress fibers in each image was calculated.

### TM phagocytosis assay

TM phagocytosis was determined by flow cytometry. Three wells were treated with control media and six wells were treated with pigment media. After 24 hours, three of the pigment treated wells were additionally treated with 10μM K-115 for 24h. After washing with PBS three times, cells were incubated with 0.5 μm FITC-labeled microspheres (F8813, Thermo Fisher, Waltham, MA) at a concentration of 5 × 10^8^ microspheres/ml for 1 h at 37°C. Wells were rinsed with PBS three times, cells digested with trypsin, resuspended in 200μl PBS, and filtered with a 70-μm cell strainer. Flow cytometry was performed to evaluate the percentage of cells that had phagocytosed FITC-labeled microspheres.

### Wound-healing assay

The ability of TM cells to migrate was assessed with a wound-healing assay. Nine wells were seeded, allowed to reach 100% confluency, and then treated with 10 μg/ml mitomycin (M4287, Sigma Aldrich)for 1 hour to inhibit cell division. Wells were washed with PBS three times and a cell-free gap was created with a single pass of a 10-μl pipette tip (F1732031, Gilson, Middleton, WI). Floating cells were gently washed away with pre-warmed media. Three wells were treated with pigment media, three wells were treated with pigment media supplemented with 10 μM K-115, and three wells were treated with vehicle control. Cells were cultured in a microscope stage incubator (H301-TC1-HMTC, Okolab, S.r.L., Ottaviano, NA, Italy), and images were taken at 40x magnification, every 6 h for 48 h using a live cell microscope system (Nikon Eclipse TI-E, Nikon, Tokyo, Japan). Cells that migrated into the cell-free gap were counted in 10 high-power fields.

### Statistics

All quantitative data were described as the mean ± standard error (SE) and analyzed by PASW 18.0 (SPSS Inc., Chicago, IL, USA). A p-value ≤ 0.05 was considered statistically significant. We used a paired t-test to compare the IOP changes from baseline and one-way ANOVA to compare several groups.

## Results

### IOP reduction

Baseline IOPs of the three groups of eyes were very similar after48 hours of stabilization (C: 9.5 ± 1.8 mmHg, n = 8; P: 11.3 ± 0.6 mmHg, n = 8; PK115: 10.0 ± 1.1 mmHg, n = 7, P = 0.56). After pigment treatment for 48 hours, IOP increased in P by 114% to 17.8 ± 1.6 mmHg (compared to C: 8.3 ± 1.8 mmHg, P <0.001). Before adding K-115, IOP increased in PK115 by 65% to 13.7 ± 1.5 mmHg (compared to C, P <0.01). There was no difference between P and PK115 (P = 0.12). IOP of C remained unchanged (P = 0.65). After 48 hours of K-115, IOP in PK115 had dropped compared to P (PK115: 8.9 ± 1.7 mmHg, P: 20.3 ± 1.2 mmHg, P <0.005). IOP of C and PK-115 were similar (11.2 ± 3.3 mmHg, n = 8, P = 0.48). One eye in the PK115 group was excluded due to contamination.

**Fig 1.**
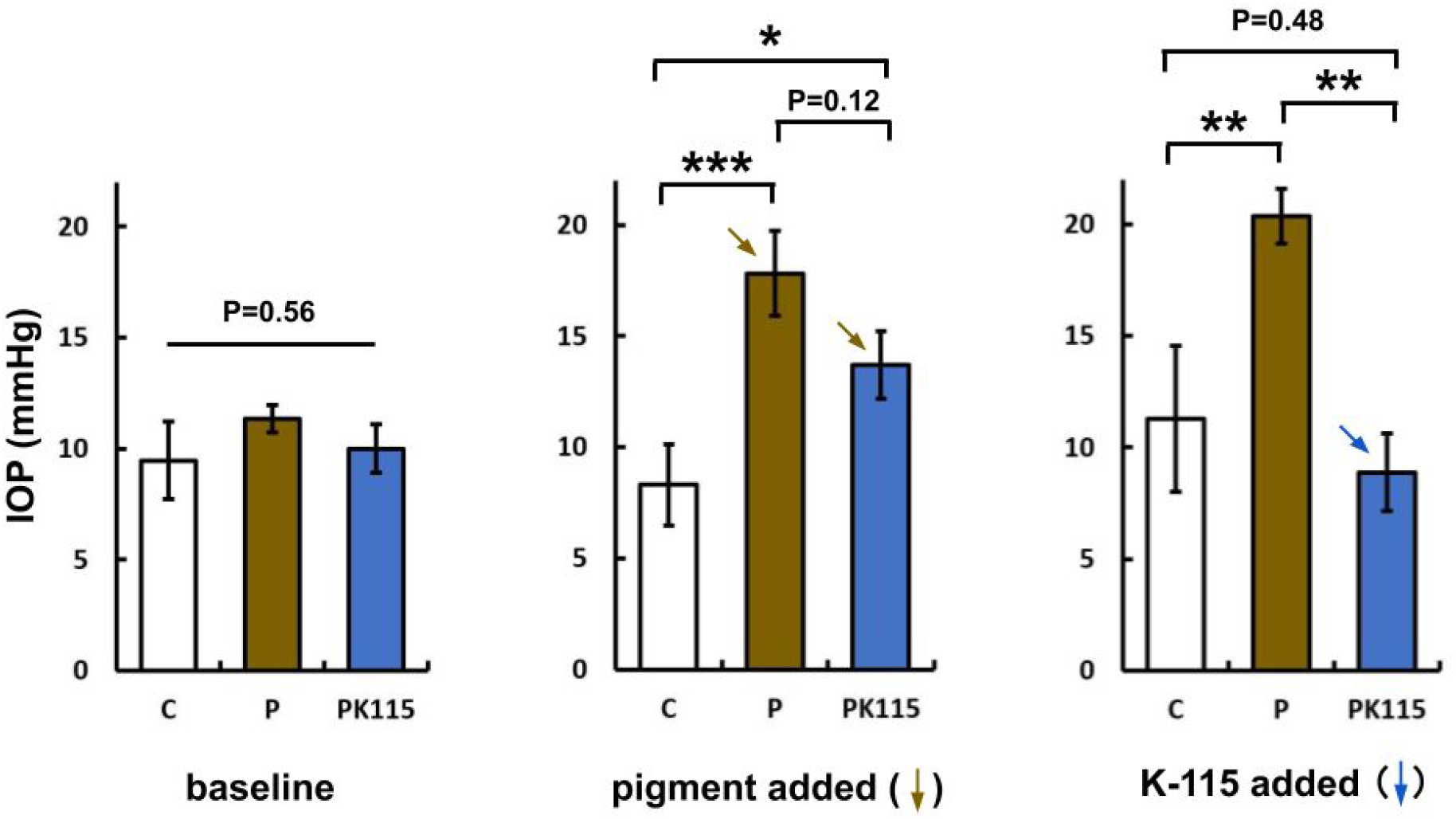
Baseline IOP of C (black), P (brown) and PK115 (blue) were similar (P = 0.56), rose in P (***P<0.001) and PK115 (*P<0.01) after pigment was added to the perfusion. After starting K-115 in PK115, IOP dropped to the level of control (difference C and PK115: P = 0.48).

### TM morphology

Three markers expressed in TM cells were selected for immunohistochemical analysis of TM primary culture: alpha-SMA, AQP1, and MGP. TM cells were positive for all three TM markers. Primary TM cells were treated with pigment media for 24 hours and observed under a phase-contrast microscope. TM cells formed a monolayer of flat and polygonal, cobblestone-like cells as expected. Pigment particles visible in P were phagocytosed and accumulated in the cells (yellow arrows). After treatment with K-115, TM cell bodies contracted and formed long, thin processes (red arrows).

**Fig 2.**
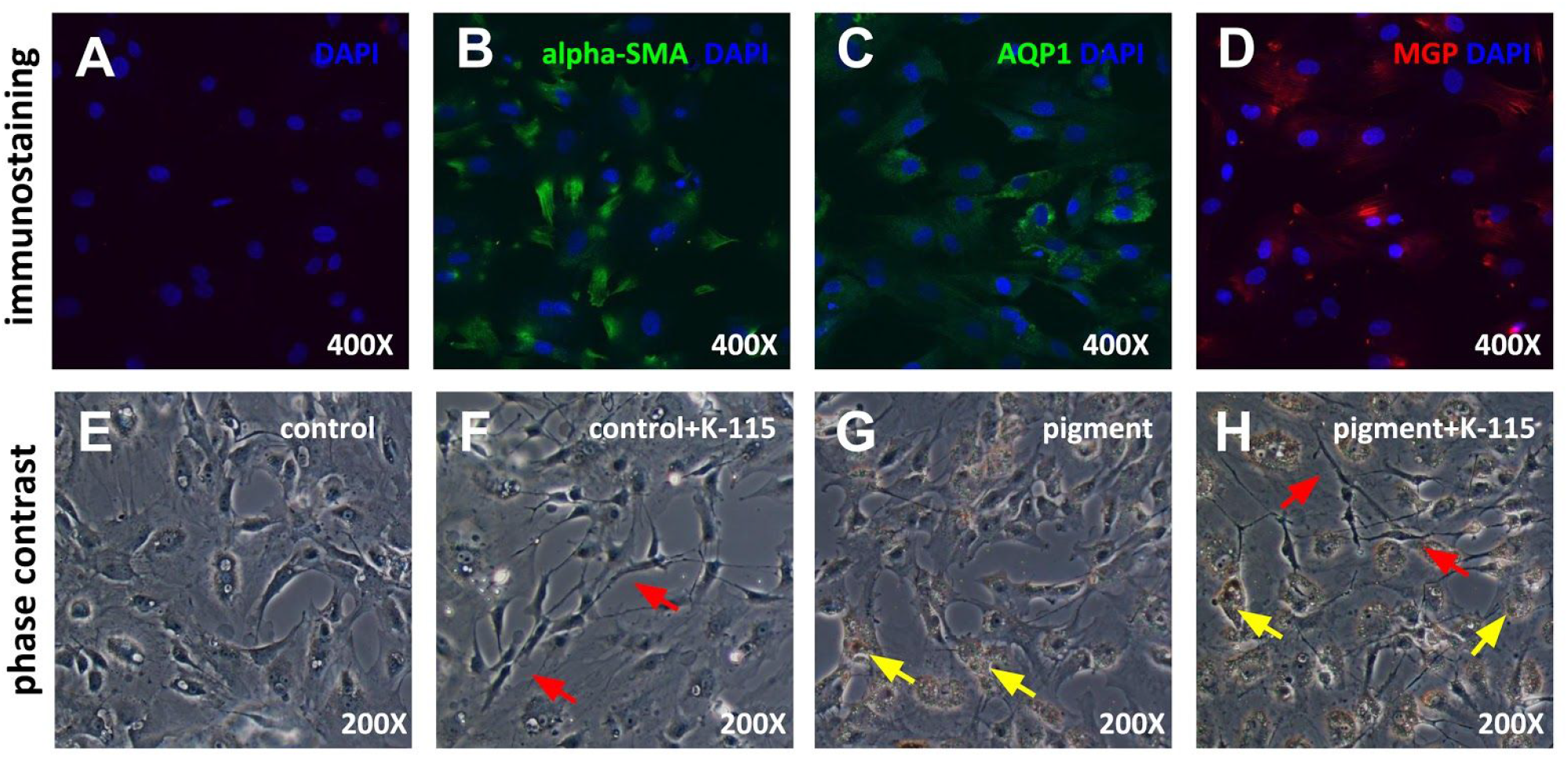
Primary TM cells expressed the TM markers alpha-SMA (B), AQP1 (C), and MGP (D). Compared to control (E), cells treated with K-115 at 10 μM contracted (F, H, red arrow). Pigment granules were phagocytosed by TM cells (G, H, yellow arrows).

### Disruption of actin stress fibers

F-actin was labeled with Alexa Fluor 488 conjugated phalloidin. In C, F-actin was thin and smooth (red arrow). After 48h of pigment treatment, F-actin in P showed thick, long, and curved stress fibers (yellow arrows). The percentage of cells that contained stress fibers in P (32.8 ± 2.9%, n = 10) was higher than that in C (21.9 ± 3.2%, n = 10, P <0.05). After 24h of pigment treatment, PK115 was treated with 10 μM K-115 for another 24 h. After treatment with K-115, the percentage of cells with stress fibers was reduced (P: 32.8 ± 2.9%; PK115: 11.6 ± 3.3%, n = 10, P <0.001).

**Fig 3.**
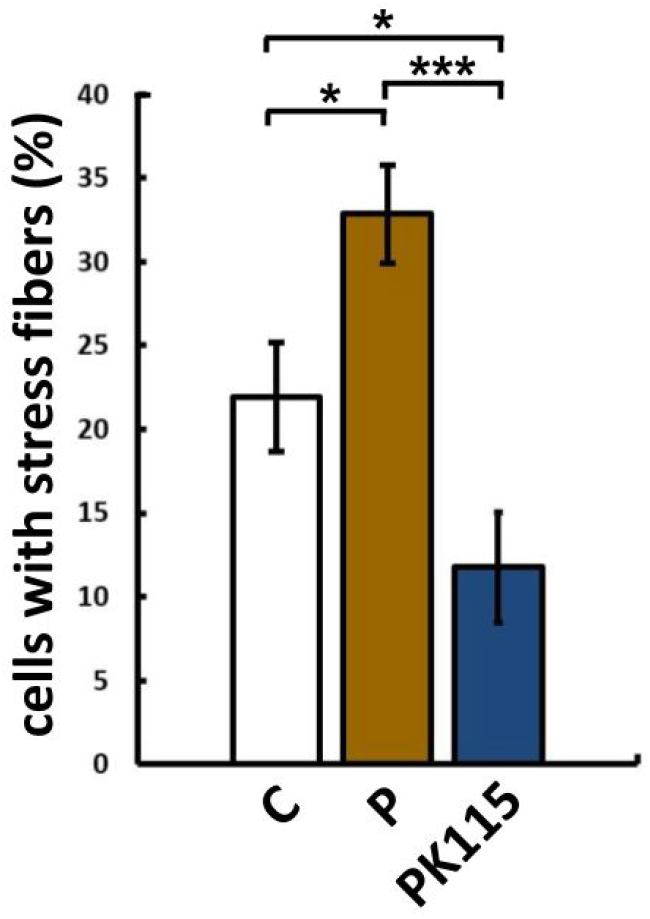
Compared with C (21.9 ± 3.2%, n = 10, * P <0.05), P (32.8 ± 2.9%, n = 10, * P <0.05) had a higher proportion of cells containing stress fibers. After K-115 treatment, TM cells containing stress fibers were reduced (C, white arrows, P: 32.8 ± 2.9%; PK115: 11.6 ± 3.3%, n = 10, *** P <0.001).

### Increased phagocytosis

After 48 h of pigment treatment, the proportion of TM cells that phagocytosed fluorescent microspheres in P was lower than C (C: 34.9 ± 0.8%, P: 25.7 ± 2.1%, n=3, P <0.01). Following 24 h of pigment treatment, PK115 was treated with 10 μM K-115 for 24 hours. The proportion of PK115 TM cells that phagocytosed fluorescent microspheres was higher than P and not different from C (P: 25.7.0 ± 2.1%, PK115: 33.4 ± 0.8%, n=3, P <0.01).

**Fig 4.**
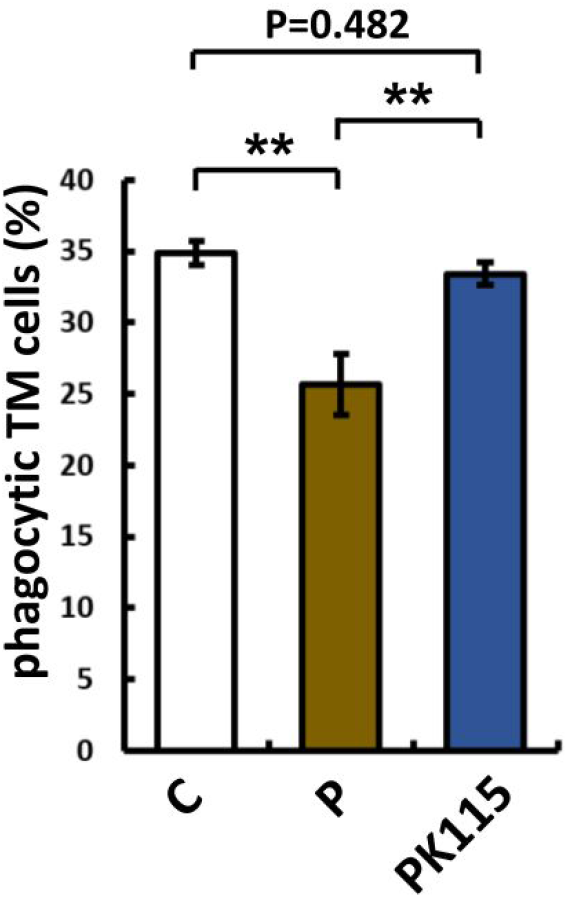
After 48 hours of pigment treatment, the proportion of TM cells that contained fluorescent microspheres in P was lower than that in C (C: 34.9 ± 0.8%, P: 25.7 ± 2.1%, n=3, ** P <0.01). After 24 hours of K-115 treatment in pigment-treated cells, the proportion of cells that phagocytosed fluorescent microspheres was higher than that in P (P: 25.7.0 ± 2.1%, PK115: 33.4 ± 0.8%, n=3, ** P <0.01).

### Increased cell motility

A wound-healing experiment was used to assess the migratory capacity of TM cells. The number of TM cells that migrated into the cell-free wound was counted at 12h, 24h, and 36h after pigment treatment. TM cells gradually migrated into the cell-free zone at 12h, 24h, and 36h after pigment treatment. TM cells had lower migration in P than in C at all time points (12h: C: 41.5 ± 4.6, P: 19.1 ± 4.6, P <0.001; 24h: C: 77.9 ± 5.2, P: 44.6 ± 4.0, P <0.001; 36h: C: 114.0 ± 6.1, P: 76.0 ± 6.9, n = 10, P <0.001). Compared with P, PK115 had higher levels of migration into the wound site at each time point as well (12h: P: 19.1 ± 4.6, PK115: 42.5 ± 1.6, P <0.001 24h: P: 44.6 ± 4.0, PK115: 64.1 ± 2.3, P <0.001; 36h: P: 76.0 ± 6.9, PK115: 100.9 ± 3.7, n = 10, P <0.001, Figure 3-3).

**Fig 5.**
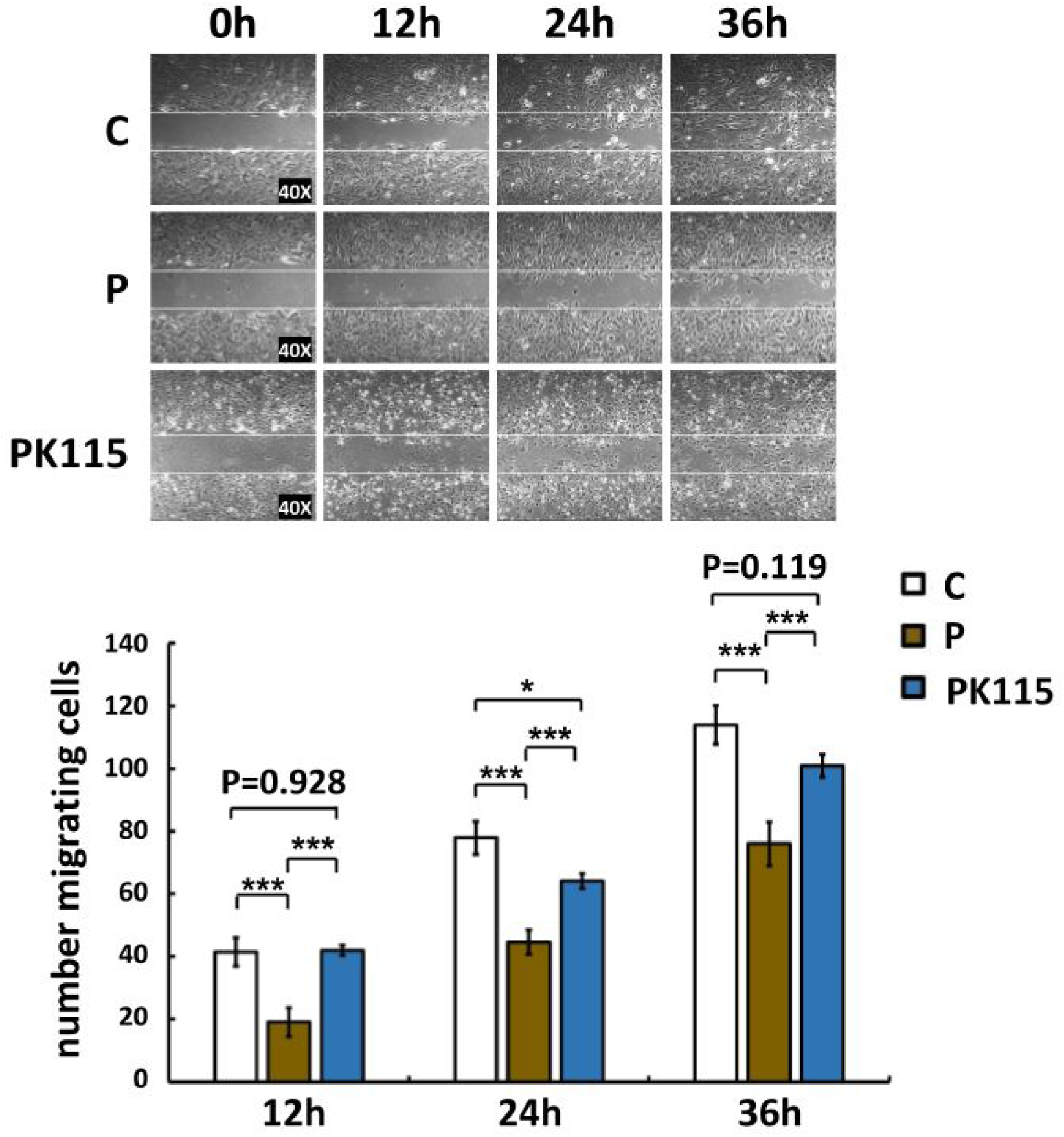
The number of TM cells that migrated to the cell-free wound in P at 12 hours (C: 41.5 ± 4.6, P: 19.1 ± 4.6, n = 10, *** P <0.001), 24 hours (C: 77.9 ± 5.2), P: 44.6 ± 4.0, n = 10, *** P <0.001) and 36 hours (C: 114.0 ± 6.1, P: 76.0 ± 6.9, n = 10, *** P <0.001,) was less than C; and K-115 increased the number of TM cells migrating to the cell-free zone at 12h (P: 19.1 ± 4.6, PK115: 42.5 ± 1.6, n = 10, *** P <0.001); 24h (P: 44.6 ± 4.0, PK115: 64.1 ± 2.3, n = 10, *** P <0.001); 36h (P: 76.0 ± 6.9, PK115: 100.9 ± 3.7, n = 10, *** P <0.001).

## Discussion

Pigmentary glaucoma is caused by the dispersion of pigment from the posterior side of the iris and ciliary processes that accumulates in the TM. Dysfunction of TM leads to increased outflow resistance and elevated IOP. In earlier studies with primary porcine TM cells [3] and porcine anterior segment perfusion model [4], we found that pigmentary debris caused an increase of TM cell stress fibers, reduced phagocytosis, and mobility, and led to a persistent IOP elevation [17]. Through gene microarray and signal pathway analysis, we found that the Rho pathway was activated in TM cells [3, 4]. Here, we investigated whether the Rho kinase inhibitor K-115 would reverse some of these changes both in vitro and ex vivo.

Several studies found that Rho-kinase inhibitors Y-27632, Y-39983, HA-1077, and H-1152, could increase outflow and decrease IOP by altering the TM cytoskeleton [8, 18–21]. Stress fibers can contract [22] and increase stiffness [23, 24] leading to decreased outflow in primary open-angle glaucoma [25] and also in glucocorticoid-induced glaucoma [26]. In our study, pigment dispersion caused a marked increase in stress fibers and IOP but then dropped to baseline after K-115 treatment.

The TM removes debris by phagocytosis [27] and has the ability to present antigens to the immune system [28]. It has been observed that porcine TM cells phagocytose pigment granules within a few hours. When the pigment concentration exceeds a certain threshold, TM phagocytosis will decrease [3, 4]. Similarly, this study showed that phagocytosis of TM cells decreases after pigment treatment and but that K-115 promotes fluorescent microsphere intake again.

Several studies found that reduced TM cell mobility may cause IOP elevation. Fujimoto et al. reported that dexamethasone-induced cross-linked actin networks (CLANs) and decreased cell migration [29]. Y-27632 counteracted the formation of CLANs. Clark found that Y-27632 lowered IOP and promoted TM cell migration in rabbits [9] similar to our observation that K-115 recovered TM mobility and IOP. It increased endothelial cell migration [30] above normal, but in our study, did not change TM cell migration above baseline.

We have previously examined RKI-1447 in the model used here. Although RKI-1447 is a pyridyl thiazolyl-urea structurally different from K-115, an isoquinoline, it is also a type I kinase inhibitor that binds the ATP binding site through interactions with the hinge region and the DFG motif and inhibits both ROCK1 and ROCK2 [31]. It has a half-maximal inhibitory concentration (IC_50_) of 14.5 nM and 6.2 nM for ROCK1 and ROCK2, respectively. In our studies, RKI-1447 achieved an IOP reduction of 33.6%. Although the present study suggests a larger IOP reduction of 50% for K-115, a side by side comparison to RKI-1447 would be helpful. RKI-1447 and K-115 have a similar specificity for ROCK 1 over ROCK2, but RK-1447 has no effect on the phosphorylation levels of AKT, MEK, and S6 kinase even at high concentrations.

In conclusion, K-115, a ROCK inhibitor, decreased TM stress fibers, increased phagocytosis, and motility in vitro, and lowered IOP ex vivo in a model of PG.

## References

1. Okafor K, Vinod K, Gedde SJ (2017) Update on pigment dispersion syndrome and pigmentary glaucoma. Curr Opin Ophthalmol 28:154–160

2. Schmetterer L (2014) Vascular optic neuropathies versus glaucomatous optic neuropathy. Acta Ophthalmologica 92:0–0

3. Wang C, Dang Y, Loewen RT, et al (2019) Impact of pigment dispersion on trabecular meshwork cells. Graefes Arch Clin Exp Ophthalmol. https://doi.org/10.1007/s00417-019-04300-7

4. Dang Y, Waxman S, Wang C, et al (2018) A porcine ex vivo model of pigmentary glaucoma. Sci Rep 8:5468

5. Mastropasqua L, Carpineto P, Ciancaglini M, Gallenga PE (1999) A 12-month, randomized, double-masked study comparing latanoprost with timolol in pigmentary glaucoma1. Ophthalmology 106:550–555

6. Grierson I, Jonsson M, Cracknell K (2004) Latanoprost and pigmentation. Jpn J Ophthalmol

7. Niyadurupola N, Broadway DC (2008) Pigment dispersion syndrome and pigmentary glaucoma–a major review. Clinical & experimental

8. Honjo M, Tanihara H, Inatani M, et al (2001) Effects of rho-associated protein kinase inhibitor Y-27632 on intraocular pressure and outflow facility. Invest Ophthalmol Vis Sci 42:137–144

9. Koga T, Koga T, Awai M, et al (2006) Rho-associated protein kinase inhibitor, Y-27632, induces alterations in adhesion, contraction and motility in cultured human trabecular meshwork cells. Exp Eye Res 82:362–370

10. Dang Y, Wang C, Shah P, et al (2019) RKI-1447, a Rho kinase inhibitor, causes ocular hypotension, actin stress fiber disruption, and increased phagocytosis. Graefes Arch Clin Exp Ophthalmol 257:101–109

11. Rao PV, Deng P-F, Kumar J, Epstein DL (2001) Modulation of aqueous humor outflow facility by the Rho kinase--specific inhibitor Y-27632. Invest Ophthalmol Vis Sci 42:1029–1037

12. Chen S, Waxman S, Wang C, et al (2020) Dose-dependent effects of netarsudil, a Rho-kinase inhibitor, on the distal outflow tract. Graefes Arch Clin Exp Ophthalmol. https://doi.org/10.1007/s00417-020-04691-y

13. Inoue T, Tanihara H (2017) Ripasudil hydrochloride hydrate: targeting Rho kinase in the treatment of glaucoma. Expert Opin Pharmacother 18:1669–1673

14. Wei L, Surma M, Shi S, et al (2016) Novel Insights into the Roles of Rho Kinase in Cancer. Arch Immunol Ther Exp 64:259–278

15. Yamaguchi H, Kasa M, Amano M, et al (2006) Molecular mechanism for the regulation of rho-kinase by dimerization and its inhibition by fasudil. Structure 14:589–600

16. Tanihara H, Inoue T, Yamamoto T (2016) One-year clinical evaluation of 0.4% ripasudil (K-115) in patients with open-angle glaucoma and ocular hypertension. Acta

17. Wang C, Dang Y, Shah P, et al (2020) Intraocular pressure reduction in a pigmentary glaucoma model by Goniotome Ab interno trabeculectomy. PLoS One e0231360

18. Tanihara H, Inatani M, Honjo M, et al (2008) Intraocular pressure-lowering effects and safety of topical administration of a selective ROCK inhibitor, SNJ-1656, in healthy volunteers. Arch Ophthalmol 126:309–315

19. Honjo M, Inatani M, Kido N, et al (2001) Effects of protein kinase inhibitor, HA1077, on intraocular pressure and outflow facility in rabbit eyes. Arch Ophthalmol 119:1171–1178

20. Fukunaga T, Ikesugi K, Nishio M, et al (2009) The effect of the Rho-associated protein kinase inhibitor, HA-1077, in the rabbit ocular hypertension model induced by water loading. Curr Eye Res 34:42–47

21. Tokushige H, Inatani M, Nemoto S, et al (2007) Effects of topical administration of y-39983, a selective rho-associated protein kinase inhibitor, on ocular tissues in rabbits and monkeys. Invest Ophthalmol Vis Sci 48:3216–3222

22. Murphy KC, Morgan JT, Wood JA, et al (2014) The formation of cortical actin arrays in human trabecular meshwork cells in response to cytoskeletal disruption. Exp Cell Res 328:164–171

23. Li G, Lee C, Agrahari V, et al (2019) In vivo measurement of trabecular meshwork stiffness in a corticosteroid-induced ocular hypertensive mouse model. Proc Natl Acad Sci U S A 116:1714–1722

24. Wang K, Read AT, Sulchek T, Ethier CR (2017) Trabecular meshwork stiffness in glaucoma. Exp Eye Res 158:3–12

25. Hoare M-J, Grierson I, Brotchie D, et al (2009) Cross-linked actin networks (CLANs) in the trabecular meshwork of the normal and glaucomatous human eye in situ. Invest Ophthalmol Vis Sci 50:1255–1263

26. Yuan Y, Call MK, Yuan Y, et al (2013) Dexamethasone induces cross-linked actin networks in trabecular meshwork cells through noncanonical wnt signaling. Invest Ophthalmol Vis Sci 54:6502–6509

27. Gagen D, Filla MS, Clark R, et al (2013) Activated *α* v *β* 3 integrin regulates *α* v *β* 5 integrin-mediated phagocytosis in trabecular meshwork cells. Invest Ophthalmol Vis Sci 54:5000–5011

28. Taylor AW (2009) Ocular immune privilege. Eye 23:1885–1889

29. Fujimoto T, Inoue T, Inoue-Mochita M, Tanihara H (2016) Live cell imaging of actin dynamics in dexamethasone-treated porcine trabecular meshwork cells. Exp Eye Res 145:393–400

30. Okumura N, Inoue R, Okazaki Y, et al (2015) Effect of the Rho Kinase Inhibitor Y-27632 on Corneal Endothelial Wound Healing. Investigative Opthalmology & Visual Science 56:6067

31. Patel RA, Forinash KD, Pireddu R, et al (2012) RKI-1447 is a potent inhibitor of the Rho-associated ROCK kinases with anti-invasive and antitumor activities in breast cancer. Cancer Res 72:5025–5034

